# Effects of matric versus osmotic potential changes on *Variovorax beijingensis* transcription

**DOI:** 10.1101/2025.06.20.660813

**Authors:** Jiwoo Kim, Bjorn Shockey, Kirsten S. Hofmockel, Xiaodong Gao, Caroline A. Masiello, Jonathan J. Silberg

## Abstract

Soil microbes must continuously adapt to changes in water availability, which dynamically fluctuates with weather and irrigation, and these adaptations are closely linked to soil CO_2_ emissions. Soil water potential, which regulates microbe-available water, is controlled by both osmotic and matric potential, which both increase as soils dry. While both parameters can independently increase water potential, the genetic mechanisms underlying microbial responses to both are unknown, with potentially different mechanisms available for soil microbes to respond to these hydrologic parameters. To explore microbial responses to matric versus osmotic potential shifts, we evaluated the growth and transcription of *Variovorax beijingensis* in soils and liquid cultures of varying water potential. We find this microbe respires in dilute minimal medium (-240 ±104 kPa), in liquid medium supplemented with sucrose (-1323 ±20.8 kPa), and in a pair of matrices that span a similar range of pressures (-183 ±55 and -1393 kPa ±200 kPa). We show that the global gene expression patterns vary significantly across all four conditions, even when the matric potential and osmotic pressure are set to similar values. However, the direction of gene expression changes correlated for 68% of the transcripts arising from an increase in osmotic pressure within liquid medium and an increase in matric potential within the different soils. While a large overlap was observed in the *Variovorax* transcriptional response to shifts in both osmotic and matric potential, the responses were not identical, with matric potential shifts leading to 2.55-fold more genes exhibiting differential expression.

**IMPORTANCE:** It remains hard to establish how changes in soil water properties affect microbial behaviors that regulate soil health, and the energy with which soil water is held is likely a holistic control on at least some of those microbial behaviors. This energy is controlled by parameters associated with soil saltiness (osmotic potential) and texture (matric potential), which both alter bioavailable water by contributing to total soil water potential. To investigate how the global transcriptional profile of a soil microbe changes when the microbe-available water is altered either by changing soil texture or by changing osmolyte concentrations, we varied osmotic and matric potential individually and performed RNA sequencing. We observe differences in the transcriptome across all conditions analyzed. A larger number of genes are differentially expressed as matric potential increases; however, many of the transcripts differentially expressed as osmotic pressure increases covary with those observed as the matrix potential increases.

## INTRODUCTION

Soil microbes live in a dynamic environment where hydrologic extremes are intensifying with climate change (1–6). Wet-dry cycles can trigger increased CO_2_ emissions through spikes of respiration that occur following soil rewetting (7–11). In part, this increase in metabolic activity occurs because microbial growth is stimulated upon soil hydration (12). This hydration response is a well-known, but incompletely constrained driver of soil CO_2_ release to the atmosphere. Microbial gene expression also varies during the growth phases following hydration (13). Following hydration, soil microbes initially focus on generating energy and repairing DNA, and then adjust their gene expression to acquire carbon and energy (13). Soil studies have also revealed that characteristics of the extracellular environment contribute to the spike in respiration following hydration, including relatively stable characteristics like soil texture and environmental parameters that dynamically change during dry-wet cycles (14), such as the concentration of osmolytes (15), carbon and nitrogen metabolite concentrations (16), and the soil matric potential (17), a property driven by capillary forces. Our understanding of drivers of respiration changes following soil wet-dry cycles remain limited because in the natural environment, multiple abiotic and biotic parameters simultaneously vary.

The amount of bioavailable water in soil is controlled by soil water potential (4′) (18, 19), which affects microbial growth and activity following rewetting (20). The total soil water potential (Ϋ_total_) is determined by four parameters (Ϋ_total_ = 4′_o_ + 4′_m_ + 4′_g_ + 4′_p_), including osmotic potential (4′_o_), matric potential (4′_m_), gravity potential (4′_g_), and the hydrostatic pressure of the water column (4′_p_). A recent study highlighted technical gaps that limit our understanding of biological responses to changes in soil water potential (18).

*In situ* measurements of water potential are limited (18), and they are rarely performed in parallel with multi-omics measurements (21). This gap constrains the data available to study the mechanistic responses of soil microbes to dynamic moisture changes in field studies. Osmotic and matric potential covary with dry-wet cycles in soil since hydration decreases the concentration of dissolved solutes and the interaction of water molecules with soil particles. This makes it challenging to disentangle the individual effects of osmotic and matric potential on microbial behaviors.

Microbial responses to osmotic stress have been intensively studied (22). These responses include synthesizing compatible solutes like proline, trehalose, and betaine (23–25), taking up solutes from the environment (26), and secreting polysaccharides (27, 28). The ecohydrological role of soil texture on plant fitness has studied in the context of rainfall fluxtuations (29), and microbial responses to textured environments have also been investigated. With some microbes, cell viability decreases with increases in matric pressure arising from soil drying (30, 31), with some microbes presenting a greater sensitivity to matric potential shifts (32), as observed with osmotic stress. Further, studies examining microbial behaviors have revealed that both the matrix and hydration level influence phenotypes (33). Some studies have compared microbial responses to osmotic and matric potential shifts. In one study, the concentrations of microbial metabolites changed more due to increases in osmotic pressure compared to matric potential (34). In another study, increased matric potential was more detrimental to microbial respiration than a corresponding change in osmotic potential (35). While these studies suggest that microbes may cope with variations in water potential arising from osmotic and matrix stress in different ways, the effects of these different types of stress on global gene expression have not been rigorously compared for individual soil microbes.

To better understand how soil microbes respond to osmotic and matric potential changes of similar magnitudes, we varied these parameters individually and studied their effect on the plant-growth promoting microbe *Variovorax beijingensis* (36). We targeted a strain from a model soil community that was previously found to support chitin degradation (37). By monitoring respiration, we find that *V. beijingensis* persists across a wide range of soil water potentials when altered by either adding osmolytes or changing soil texture (altering matric potential). Using RNA sequencing (38), we map the transcriptional response to increases in soil water potential arising from both osmotic and matrix stress. While this analysis revealed significant differences in the transcriptional responses upon osmotic and matrix stress, a large fraction (68%) of the genes covaried in their differential expression upon increases in osmotic and matric potential.

## MATERIALS AND METHODS

### Cell Growth

*V. beijingensis*, which has the strain designation PNNL_MSC-1, was isolated from a field site in Prosser, WA (37). This strain was grown in a variety of media, including Luria-Bertani (LB) medium, M9 minimal medium, and modified M9 medium (mM9). LB (per liter) was prepared by mixing tryptone (10 grams), NaCl (5 grams), and yeast extract (5 grams). M9 contained M9 salts, glucose (0.4%), MgSO_4_ (2 mM), and CaCl_2_ (0.1 mM). M9 salts were prepared as a 5x stock by mixing KH_2_PO_4_ (15 grams), Na_2_HPO_4_·7H_2_O (64 grams) NaCl (2.5 grams) and NH_4_Cl (5 grams) per liter of water. The modified M9 (mM9) was designed to make liquid media with lower water potential while keeping the concentration of the glucose and NH_4_Cl the same concentration as M9 medium. The 5x salt stock was prepared identically, except no NH_4_Cl was included. Instead, it was added at a concentration of 1 gram per liter in the mM9. To adjust the water potential of mM9 media, two approaches were used. Either the concentration of salt stock added was varied, or sucrose was added. LB medium, 5x M9 and mM9 salt stocks were all autoclaved, while other growth medium reagents were sterile filtered through a 0.2 µm filter (Thermo Scientific and Satorius AG). Chemicals for making growth media were from Sigma-Aldrich and Thermo Scientific.

### Artificial Soil Preparation

Artificial soils were produced using a previously described approach (31). Briefly, nonreactive quartz materials of three different particles sizes (whole grain fine quartz sand with particle size ∼70 μm, ground silt-sized quartz with particle size ∼8.71 μm and clay-sized quartz with particle size ∼1.7 μm obtained from US Silica) were used to represent soil particles of sand, silt, and clay, respectively. By varying the percentage of the three quartz particles, we produced artificial soils with desired soil textures. In addition to soil texture, soil mineralogy also affects soil matric potential (39). Therefore, two common clay minerals, kaolinite and montmorillonite (Spectrum Chemical MFG Corp) were also used to represent more chemically reactive clay fraction of the artificial soils. Three of these soils were quartz based and varied in texture only, including sand (90% sand, 5% silt, 5% clay), silt loam (20% sand, 60% silt, 20% clay), and clay (20% sand, 20% silt, 60% clay); these are named Quartz-1 (Q1), Quartz-2 (Q2), and Quartz-3 (Q3), respectively. Two additional silt loam soils were created which contain the clay minerals montmorillonite (M2; 20% sand, 60% silt, 20% montmorillonite) and kaolinite (K2; 20% sand, 60% silt, 20% kaolinite).

### Water Potential Analysis

The water potential of growth medium and hydrated soils was measured using WP4C dew point potentiometer (METER Group) at room temperature. The dew point potentiometer measures the total water potential of a sample and does not differentiate osmotic and matric potential. The WP4C was calibrated using a 0.5 M potassium chloride standard (AQUALAB by Decagon) to a pressure of -2.20 MPa. All measurements were made using three replicates using precision mode. The dry weight of each artificial soil was hydrated to θ = 0.1 g/g with mM9 and allowed to equilibrate overnight prior to measurement.

### Growth in Liquid Medium

Colonies of *V. beijingensis* were obtained by growing on LB-agar plates for 72 hours at 30°C. Single colonies were used to inoculate M9 medium (3 mL), which were grown at 30°C while shaking at 250 rpm for 72 hours. The resulting precultures were used as inoculants for all subsequent cultures. To obtain growth curves, the preculture was washed with each medium (1 mL) used in subsequent growth experiments three times. Cells were then resuspended in growth medium (100 µL) to optical density (OD) of 0.05 in 96-well plates (Nuclon Thermo Fisher). Cell growth was monitored continuously for 72 hours by measured the OD every 5 minutes, while shaking (2.5 mm amplitude at 105 rpm) at 30°C using a plate reader (Tecan Spark). To evaluate liquid growth containing varying concentration of M9 salts and sucrose, samples were treated identically, except they were resuspended to an OD of 0.01 in total volume of 500 µL in a 96-well deep well plate. Following incubation at 30°C, while shaking at 250 rpm, cultures (200 µL) were transferred to a transparent 96-well plate (Nuclon Thermo Fisher), and OD was measured using a plate reader. To evaluate liquid growth in vials, the preculture was washed three times with mM9 and diluted to an OD of 0.1 with total volume of 1 mL in a 2-mL glass vial (Thermo Scientific 6ACV11-1PT). Cultures were immediately crimped (Thermo Scientific 6ACC11ST1T) and then incubated statically at 30°C for 72 hours. Following incubation, the headspace CO_2_ was measured. To quantify the number of CFU, each culture was diluted serially using mM9 and an aliquot of each dilution (5 µL) was spotted on mM9-agar plates. After 72 hours at 30°C, colonies were counted. All experiments were performed using three biological replicates unless stated otherwise.

### Soil Growth Analysis

One gram of each autoclaved artificial soil was placed in GC-MS vials (2 mL). These soils were then hydrated to 10% water content by adding 100 µL of a *V. beijingensis* culture that had been generated by washing the preculture with mM9 three times and resuspending it in mM9 to OD of 1, keeping the cell number constant relative to the liquid growth experiments. Vials were crimped and inoculated for 72 hours at 30°C. Headspace CO_2_ was then measured. Following this analysis, cells were hydrated with mM9 and transferred to 15 mL conical tubes to obtain cells to spread on plates. M2 required a higher volume of liquid medium (3 mL) compared to Q2 (750 µL) as it was difficult to pipette up the slurry due to its smaller particle sizes. For each sample, a proportional amount of the slurry was serially diluted using mM9, an aliquot (5 µL) of each dilution was spotted on mM9-agar plates at different dilutions, and plates were incubated for 72 hours at 30°C. Following incubation CFU were manually counted on plates. Three biological replicates were performed for each experiment.

### Headspace Gas Analysis

To monitor CO_2_ produced by cultures in sealed vials, gas chromatography-mass spectrometry (GC-MS) used for headspace analysis following 72-hour incubations. This instrument consists of Agilent 7890B GC system, a 5977B MS, and a 7693A liquid autosampler equipped with a 100 µL syringe. Headspace gas (50 µL) was injected into DB-VRX capillary column (20 m, 0.18 mm I.D., and 1 μm film) at 50:1 split ratio, and the oven temperature was held at 45°C for 1 minute. MS analysis was performed using selected ion monitoring mode for CO_2_ (MW = 44 and 45). We used Agilent MassHunter Workstation Quantitative Analysis software to quantify the peak area of the major ions and used the minor ions as qualifiers. All CO_2_ levels are reported as relative peak area measured using three biological replicates.

### RNA Sequencing

To generate samples for RNA sequencing, four samples were set up within crimped 2 mL vials, including cells in liquid mM9 (1 mL) lacking sucrose at an OD of 0.2, cells in liquid mM9 (1 mL) containing 425 mM sucrose at an OD of 0.2, cells in Q2 (1 gram) hydrated to 10% water content with mM9 (100 µL) using cells at an OD of 2, and cells in M2 (1 gram) hydrated to 10% water content with mM9 (100 µL) using cells at an OD of 2. Thus, under each incubation condition, a similar number of cells were used to start cultures in mM9 medium, which varied in sucrose or matrix. Following incubation for 72 hours, the cells were lysed open with bead beater (Biospec Products). The total RNA then was extracted using ZymoBIOMICS RNA Miniprep Kit (Zymo Research) in accordance with the manufacturer’s instructions. The concentration and the quality of the extracted RNA were measured using Qubit 4 Fluorometer (Thermo Fisher Scientific) and spectrophotometer (DeNovix DS-11 FX+) respectively. The extracted RNA was prepared for Next Generation Sequencing using Zymo-Seq RiboFree Total RNA Library Kit (Zymo Research) following manufacturer’s instructions. The prepared library was sent to Azenta Life Sciences where the library sizes were validated with Agilent TapeStation (∼560 bp) and sequenced using NovaSeq 6000 XP S4. Six biological replicates were performed for each sample condition excluding one low osmotic potential condition where the DNA concentration of one of the six samples could not be detected using the TapeStation.

### Differential Gene Expression Analysis

The assembled reference genome and annotations were previously described. Raw paired-end FASTQ files were processed using a custom Python (v3.9.21) pipeline (https://github.com/SilbergLabRice/SeqSorcerer). The pipeline was modified to include parameters specific to this study. Quality trimming and FastQC analysis were performed using Trim Galore v0.6.10 (https://github.com/FelixKrueger/TrimGalore) with the settings "--fastqc", "--paired", "--length", "20","--clip_R2", "15", "--three_prime_clip_R1", and "15". The reference genome was indexed and aligned with HISAT2 v2.2.1 (40). The BAM files were processed with SAMtools v1.9 (41). Transcript counts were quantified via featureCounts from the Subread package v2.0.1 with the parameters, ’-a’, gtf_file, ’-o’, output_file, ’-T’, ’2’, ’-s’, ’2’, ’-Q’, ’0’, ’-t’, ’CDS’, ’-g’, ’transcript_id’, ’--minOverlap’, ’1’, ’--fracOverlap’, ’0’, ’--fracOverlapFeature’, ’0’, and ’-p’ (42).

After the raw sequencing reads were processed, genes with low total counts (≤10) were eliminated. The dataset pairs that were directly compared, e.g., liquid medium having high versus low osmotic pressure, were normalized and evaluated for differentially expressed genes (DEGs) using DESeq2 v1.44.0 in R v4.4.1 with Benjamini-Hochberg (BH) adjusted *p*-value capped at 0.05 (43). To conduct principal component analysis (PCA), the RNA-seq data was normalized using variance stabilizing transformation (“vst”) and “plotPCA” functions from DESeq2 package. PCA plot was generated via ggplot2 v3.5.1 (44), and the shaded area represents clustering of biological replicates with 95% confidence using multivariate t-distribution. The pairwise Euclidean distance was calculated with the first two principal components of each sample. Permutational multivariate analysis of variance (PERMANOVA; 999 permutations) test was performed with “adonis2” function using Euclidean distance method in vegan package v.2.7-1 (45).

### DEG Pathway Analysis

The Kyoto Encyclopedia of Genes and Genomes (KEGG) Orthology (ko) numbers assigned to each gene were extracted from the genome annotation (46). These ko numbers were organized by condition and direction of gene regulation. KEGG module enrichment analysis was performed using “enrichMKEGG” function of clusterProfiler package v4.12.6 (47). With the analysis, we referenced the “ko” organism database and used a BH adjusted *p*-value cutoff of 0.05 and *q*-value of 0.2. All RNA-seq data was visualized using ggplot2 (44).

### Data Analysis

Average and standard deviation of water potential, OD, and headspace CO_2_ were calculated in R. The *p*-value was generated from two-tailed, unpaired t-test in R. The data was visualized using ggplot2 (44).

## RESULTS

### Modulating water potential with osmolytes

*Variovorax* has been implicated as a drought-resistant genus (48, 48, 49), but it is not clear how the persistence of microbes in this genus varies with osmotic pressure. To establish a strategy to tune osmotic pressure in liquid culture and in soils, we varied the osmotic pressure of M9 minimal medium by diluting 5x M9 minimal salts to different extents. With these experiments, the nitrogen (ammonium chloride) and carbon (glucose) sources were held constant at the levels normally found in M9. *V. beijingensis* grew across all conditions analyzed, presenting OD values that increased with M9 salt concentrations (Figure 1a). Analysis of the pressures of each liquid medium revealed that this microbe grows across pressures ranging from -150 to -1500 kPa (Figure 1b); -1500 kPa is defined as the permanent wilting point (PWP), representative of the wilting point of most plants (50). These results show that M9 minimal medium whose salts have been diluted 10-fold (-240 kPa +/-104 kPa), which we call modified M9 medium (mM9), can be used as a frame of reference to study osmolyte and matrix effects on *V. beijingensis* viability. This liquid medium was used for all subsequent measurements.

**Figure 1.**
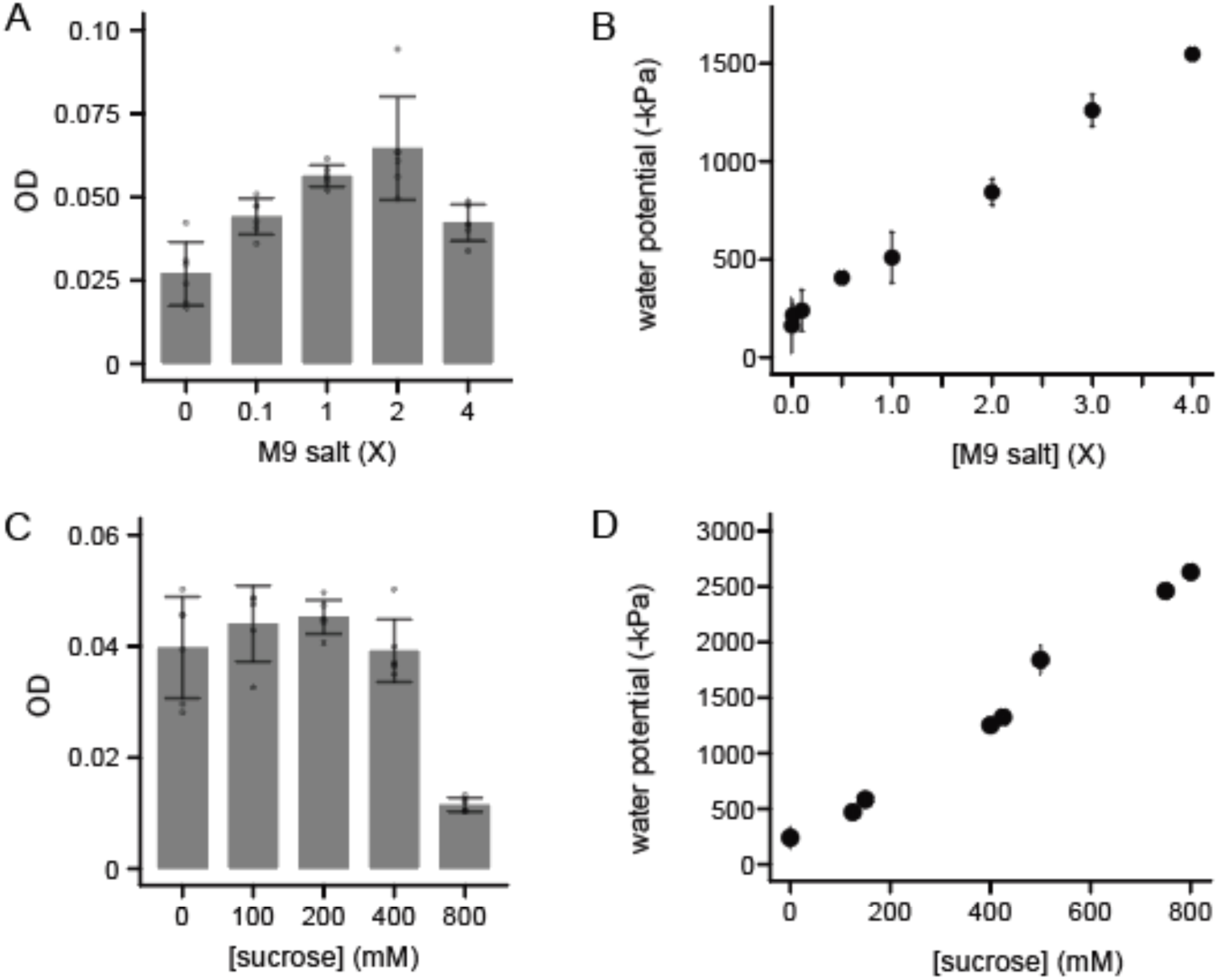
*V. beijingensis* grows across a wide range of osmotic pressures. (**A**) Optical density (OD) of *V. beijingensis* incubated aerobically for 72 hours at 30°C in M9 medium containing varying concentrations of M9 salts while shaking at 250 rpm. (**B**) Water potential of M9 medium containing different concentrations of M9 salts. (**C**) OD of 1. *V. beijingensis* grown in mM9 containing different concentrations of sucrose. Cells were incubated aerobically for 72 hours at 30°C while shaking at 250 rpm. (**D**) Water potential mM9 containing varying concentrations of sucrose. For growth measurements, the points represent biological replicates, the bars represent the average, and error bar represent ±1σ. For water potential measurements, points represent the average of three replicates, while the error bars are ±1α.

To increase the osmotic pressure of mM9 medium while minimizing effects on *V. beijingensis* metabolism, we evaluated the effect of adding sucrose on growth and osmotic pressure. This sugar was chosen because *V. beijingensis* cannot use it as carbon source (37). When cells were provided sucrose as the only carbon source in M9 medium, growth was not observed (Figure S1), while robust growth was observed with cells provided glucose. We next examined how sucrose affects the growth of *V. beijingensis* when added to mM9, which contains glucose. *V. beijingensis* grew to similar final densities in mM9 containing 0 to 400 mM sucrose (Figure 1c), although cells presented diminished growth in mM9 containing 800 mM of sucrose. To understand the pressures arising from each sucrose amendment, the potential was measured in mM9 containing different sucrose concentrations using a dew point potentiometer (Figure 1d). This analysis revealed that *V. beijingensis* grows across in mM9 having pressures ranging from -240 ±104 kPa (mM9 lacking sucrose) to -1253 ±55 kPa (400 mM sucrose). These results show that *V. beijingensis* can grow in liquid medium presenting a wide range of pressures, induced by adding an osmolyte that this microbe cannot metabolize.

To investigate if *V. beijingensis* can persist close to permanent wilting point (-1500 kPa), the total CO_2_ respired and the colony forming units (CFU) of *V. beijingensis* was measured after incubating cells statically for 72 hours at 30°C in mM9 medium lacking (-240 ±104 kPa) and containing 425 mM sucrose (-1323 ±21 kPa). The total CO_2_ accumulation in both conditions was similar (Figure 2a). Also, plating cells on mM9-agar plates following each incubation yielded similar CFU in the absence (2.6 ±2.0 x 10^9^) and presence (4.6 ±1.6 x 10^9^) of sucrose. These findings show that *V. beijingensis* can persist in mM9 medium spanning osmotic pressures that vary by >1000 kPa.

**Figure 2.**
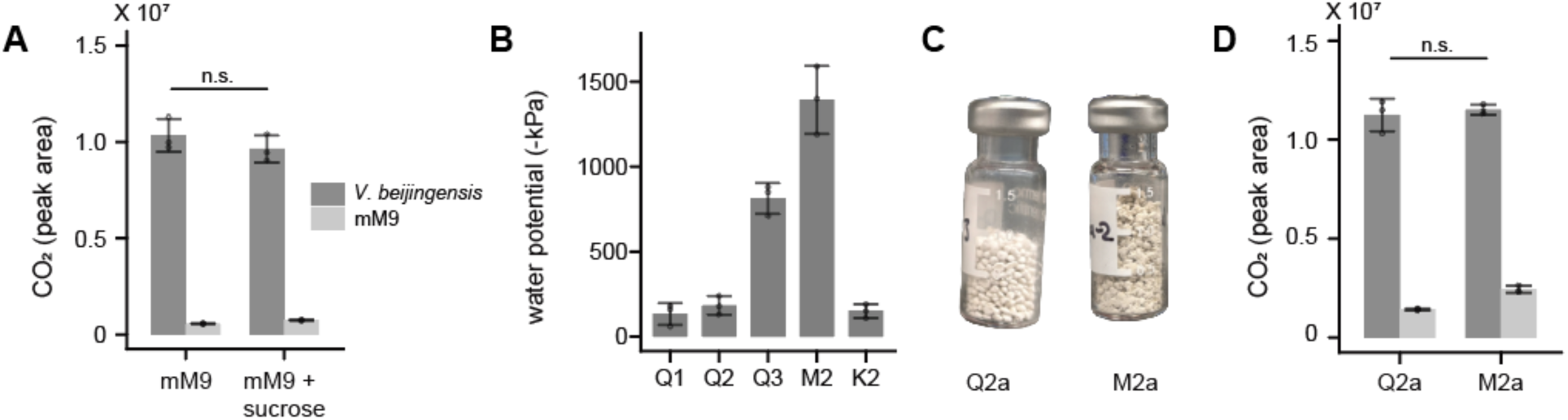
*V. beijingensis* respiration in liquid medium and artificial soils. (**A**) Headspace CO_2_ for *V. beijingensis* grown in crimped vials (*dark gray*) is compared with cultures lacking cells (*light gray*). Cells were incubated in mM9 medium containing 425 mM sucrose (-1323 kPa) or lacking sucrose (-240 kPa) for 72 hours at 30°C without shaking. (**B**) Water potential of artificial soils Q1, Q2, Q3, M2, and K2 hydrated to a water content (θ) of 0.1 gram of water per gram of matrix. (**C**) The artificial soils that were hydrated with mM9 containing *V. beijingensis* at a water content of 10% to study effects of matric potential on respiration (**D**) Headspace CO_2_ for *V. beijingensis* following 72 hours in each matrix at 30°C. Soils inoculated with cells (*dark gray*) were compared with those hydrated with mM9 lacking cells (*light gray*) for each matrix. In all experiments, points represent biological replicates, bars represent average all replicates, and error bars represent ±1α. *p*-values were calculated using two-tailed, unpaired t-test (ns, p>0.05).

### Using artificial soils to tune water potential

Artificial soils can be created that mimic soil physical properties, such as soil water energy, by varying grain size (31). To screen for artificial soils that can be used to achieve both low and high matric potential when hydrated to the same water content, we characterized the effects of adding mM9 medium on the soil water potential of five different artificial soils (Q1, Q2, Q3, M2, and K2), guided by water potential curves previously reported for these soils (31). When each soil was hydrated with mM9 medium to 100 mg of water per gram of matrix (θ = 10%), we found that Q1, Q2, and K2 yielded the lowest pressures, ranging from -133 +/- 64 kPa (Q1) to - 183 ±55 kPa (Q2) (Figure 2b). These pressures are similar to those observed with liquid mM9 medium in the absence of matrix (-240 ±104 kPa). M2 yielded the highest pressure when hydrated to 10% water content with mM9, yielding a pressure (-1393 kPa ±200 kPa) that is similar to the pressure of mM9 containing 425 mM sucrose (-1323 ±20.8 kPa). An intermediate pressure (-814 ±91 kPa) was observed with Q3. Thus, when artificial soils are hydrated to 10% water content using mM9, the M2 matrix presents a pressure that is similar to the pressure observed in mM9 containing 425 mM sucrose, while Q2 exhibits a pressure that is like the pressure of liquid mM9.

The finding that *V. beijingensis* respires and persists in mM9 medium lacking sucrose and containing 425 mM sucrose suggested that this microbe could also persist in the artificial soils presenting a similar range of pressures. To test this idea, Q2 and M2 were hydrated to 10% water content using mM9 containing *V. beijingensis* in crimped vials (Figure 2c). The number of cells used for inoculation was identical to that of the liquid medium experiments. However, cells were resuspended in smaller volumes of liquid so that they yielded the desired soil water potential following addition to the soils. After 72 hours of static incubation at 30°C, the CO_2_ produced by soils was evaluated as well as the number of CFU. *V. beijingensis* presented similar respiration in both soils (Figure 2d), which mirrored that observed with liquid cultures. In addition, cells extracted following each incubation yielded similar CFU for the Q2 (8.9 ±2.2 x 10^8^) and M2 (5.9 ±1.3 x 10^8^) soils. These findings identify soil conditions that can create water potentials that mirror the range of potentials generated by adding sucrose to liquid mM9 medium.

### Individual effects of osmotic and matrix stress on transcription

To investigate the effects of osmotic stress on *V. beijingensis* transcription, cells were grown in liquid mM9 medium lacking or containing sucrose (425 mM), total RNA was purified after a 72-hour incubation, and RNA sequencing was performed. In parallel, cells were grown in Q2 and M2 soils hydrated to 10% water content using mM9 medium containing an identical titer of *V. beijingensis*, which yields a similar range of water potentials. To determine if there are significant differences in the transcriptional profiles of cells grown across these conditions, principal component analysis (PCA) was used to compare the RNA sequencing data. Across both principal components, the separation between the liquid medium treatments presented a smaller separation than between the Q2 and M2 matrix conditions (Figure 3, Table S1). Permutational multivariate analysis of variance (PERMANOVA) using Euclidean dissimilarity revealed significant differences across all conditions and showed that 82% of the total variation is explained by the tested conditions (Table S2). These findings show that all four of the incubation conditions yield distinct patterns of global gene expression even though some of the conditions have similar water potentials.

**Figure 3.**
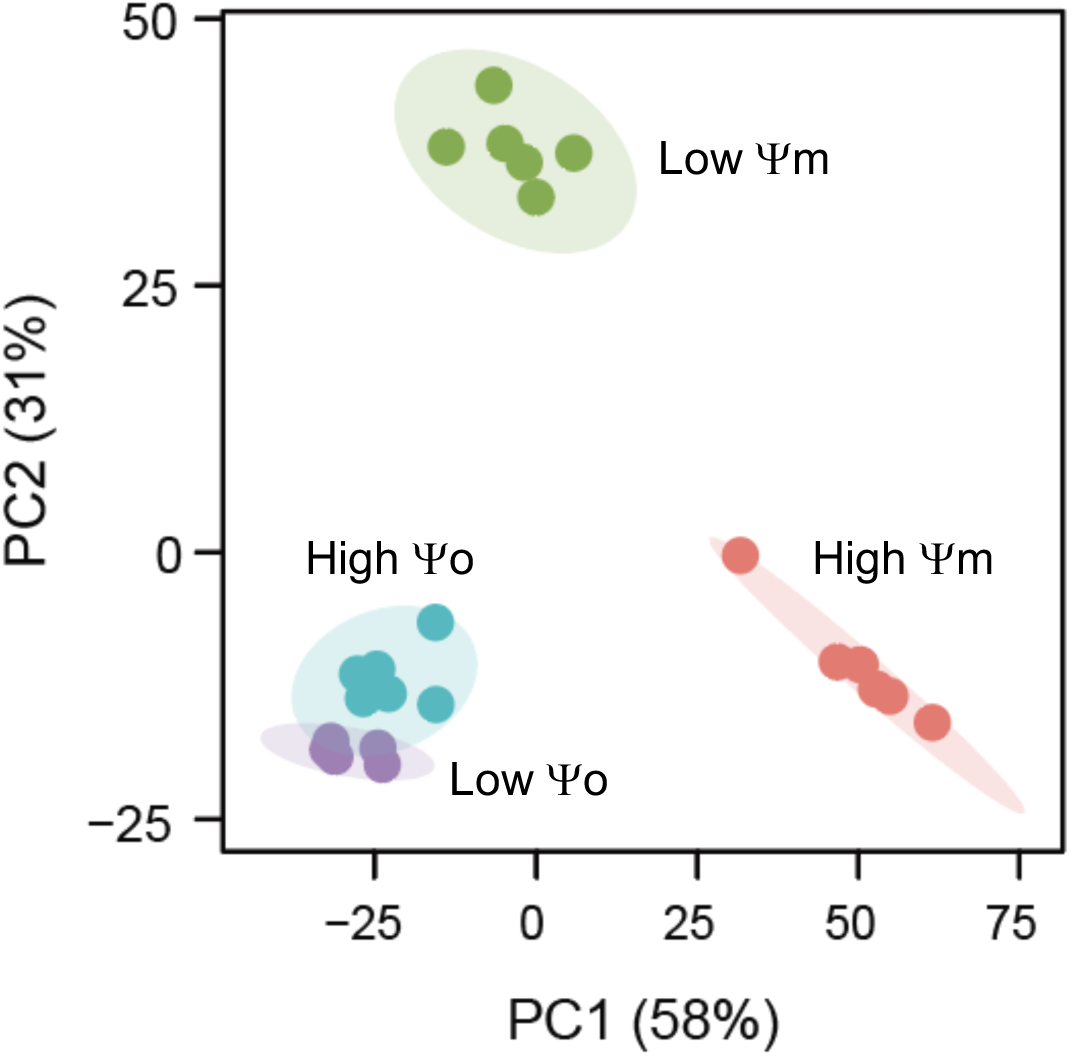
Principal component analysis of *V. beijingensis* transcription. RNA sequencing data was compared for cells grown in liquid mM9 medium lacking sucrose and having an osmotic pressure of -240 kPa (purple), liquid mM9 medium containing sucrose and having an osmotic pressure of -1323 kPa (light blue), Q2 soil hydrated to 10% water content with mM9 and having a matric potential of -183 kPa (*green*), and M2 soil hydrated to 10% water content with mM9 and having a matric potential of -1393 kPa (*red*). Each point represents a biological replicate. The shaded clusters for each set of points represent 95% confidence using multivariate t-distribution.

To understand how the gene expression patterns vary with osmotic stress in liquid medium, we evaluated how the gene expression of the 6440 annotated genes changed in expression in liquid mM9 lacking or containing sucrose (Figure 4a). Under high osmotic potential conditions (-1323 kPa), 916 genes were significantly down regulated, while 1062 genes were significantly upregulated (Wald test; p-adjusted < 0.05). To identify pathways that are differentially expressed, an enrichment analysis was performed using the KEGG to evaluate the differentially expressed genes (DEGs) presenting significant changes in expression [Benjamini-Hochberg (BH) procedure; p < 0.05, q < 0.2]. This analysis revealed that genes implicated in glucose metabolism and oxidative phosphorylation pathways are prominent among those upregulated by osmotic pressure (Figure S2), including genes involved in glycolysis, the pentose phosphate pathway, gluconeogenesis, and NADH quinone oxidoreductase. Notably, nine genes implicated in betaine metabolism were upregulated under high osmotic pressure condition, a pathway that has been implicated as an osmotic stress response in other prokaryotes (22, 51, 52). In contrast, pathways implicated in amino acid, nucleotide, and cofactor biosynthesis were down regulated under high osmotic potential (Figure S3). These experiments establish how *V. beijingensis* gene expression changes when cells experience a ∼1000 kPa increase in water potential induced by changing the osmotic pressure.

**Figure 4.**
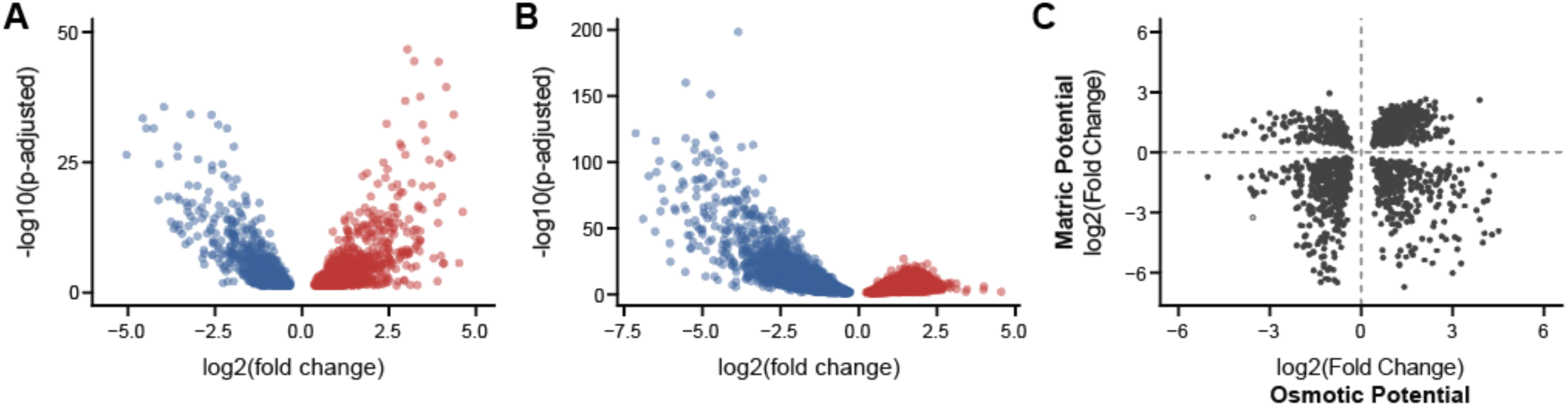
*V. beijingensis* genes that are differentially expressed genes when water potential increases. The changes in gene expression observed when comparing cells grown in (**A**) liquid mM9 medium containing or lacking sucrose, and (**B**) comparing cells grown in Q2 and M2 soils hydrated to 10% water content with mM9 medium containing cells. Blue data points represent genes that are downregulated as the soil potential increases, while red data points represent those that are upregulated as soil potential increases. (**C**) A comparison of DEGs arising from osmotic stress and matrix stress. Among all genes that showed significant changes in gene expression under both osmotic and matrix stress (n = 1508 unique genes), approximately two thirds covaried in the direction of those gene expression change (n = 1021 genes). Only those genes that presented a significant change in gene expression (Wald test; P_adjusted_ <0.05) are shown.

To establish how cells grown in soils having different matric potentials influence gene expression, we investigated how gene expression varies across Q2 and M2 matrices. Under the high-water potential growth condition of the M2 matrix, large numbers of genes were significantly upregulated (n = 2678) and downregulated (n = 2374) compared to cells grown in the low potential Q2 matrix (Figure 4b). In total, 2.55-fold more genes presented significant changes in gene expression across the two matrices compared with measurements comparing the two liquid growth conditions. When comparing gene expression in M2 to Q2, pathways having large numbers of upregulated genes included those implicated in glucose metabolism, oxidative phosphorylation, and betaine metabolism (Figure S4). Pathways having the largest numbers of genes downregulated included those implicated in the amino acid biosynthesis, nucleotide metabolism, cofactor synthesis, and the citrate cycle (Figure S5). These measurements show how gene expression changes when cells experience a ∼1000 kPa increase in water potential induced by altering the matrix, while holding water content constant.

### Comparing the effects of osmotic and matrix stress

Gene expression measurements revealed that matrix changes induce more DEGs than osmolyte changes. To better understand how these DEGs relate under the different water potential conditions evaluated, the DEGs arising from osmotic and matrix stress were compared (Figure 4c). In total, 1508 genes presented a significant change in gene expression when subjected to both matrix and osmotic stress. These DEGs had a large overlap (76%) with the genes that exhibited differential expression in liquid medium. Among these shared DEGs, 1021 genes (67.7%) covaried in their direction of differential gene expression when cells experienced both osmotic and matrix stress. Thus, although a smaller number of genes present altered gene expression under osmotic stress, a majority of these DEGs present similar responses under matrix stress.

To understand the pathways that covaried with osmotic and matric potential increases, KEGG enrichment analysis was performed. Metabolic pathways that were upregulated under both mechanisms of water potential changes included pathways implicated in betaine and amino acid metabolism (Figures 5a). Some of the most prevalent pathways that were upregulated under both types of stress were those involved in betaine metabolism and energy production, such as glycolysis, gluconeogenesis, and the pentose phosphate pathway. Also, the genes encoding 1-aminocyclopropane-1-carboxylate (ACC) deaminase and nodulation factor transporter were upregulated in both conditions; these represent plant growth promoting genes (53). Pathways involved in amino acid biosynthesis were prevalent among the DEGs that were downregulated under both types of stress (Figure 5b). A much smaller number of pathways had DEGs where the direction of the gene expression change was not correlated (Figure 5c). Among these pathways, cobalamin synthesis and KDO2-lipid A biosynthesis were upregulated by matrix stress and downregulated by osmotic stress, while leucine/lysine degradation and NADH-quinone oxidoreductase were downregulated by matrix stress and upregulated by osmotic stress. Together, these analyses show that a majority of the DEGs arising from shifts in osmotic and matric potential exhibit similar directions of expression changes. They also identify the metabolic pathways having DEGs that respond similarly to increases in water potential arising from increased osmolyte concentrations and altered soil texture that increases matric potential.

**Figure 5.**
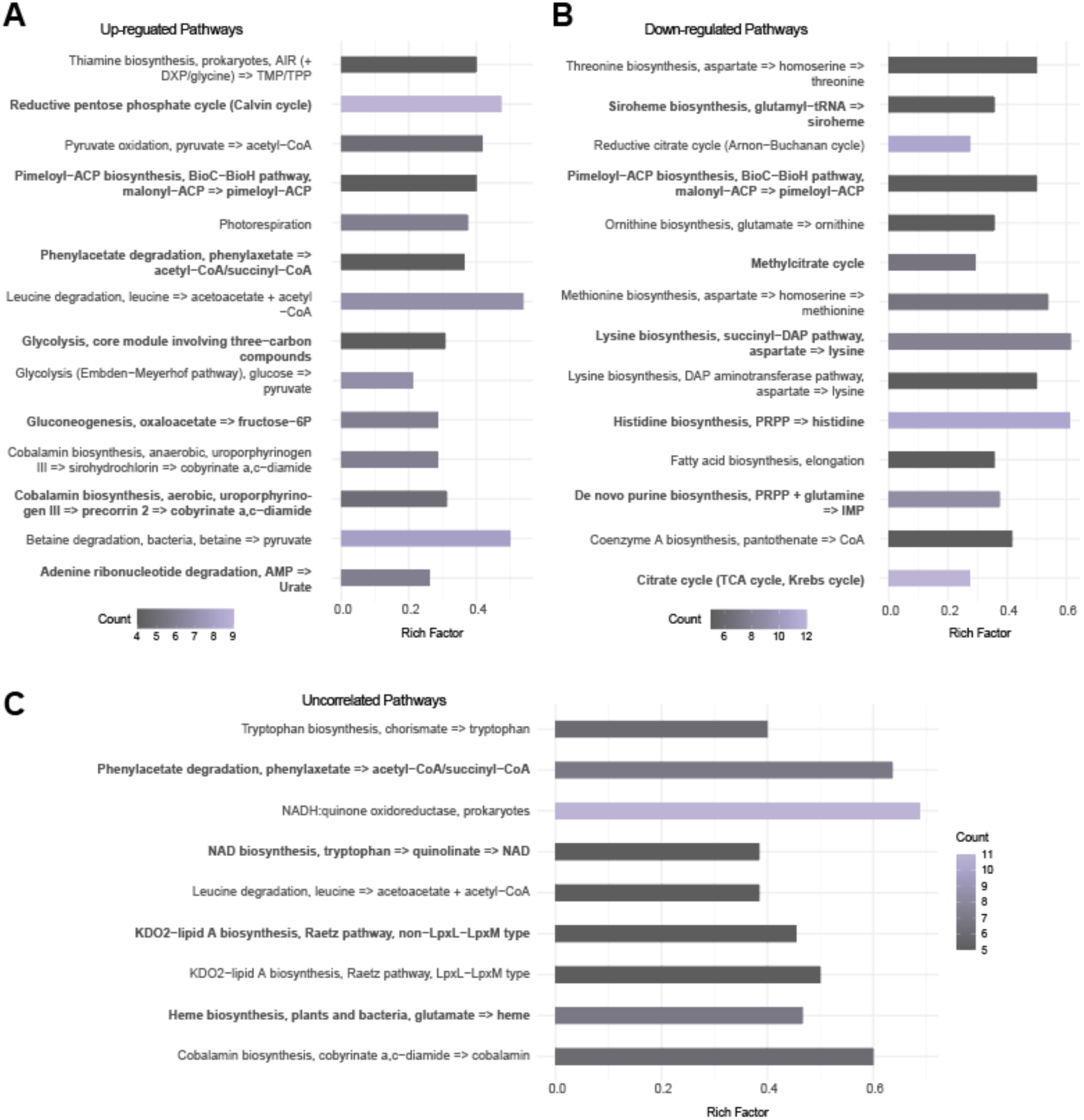
Comparisons of pathways that are differentially expressed under both matrix and osmotic stress. (**A**) DEGs upregulated under both osmotic and matrix stress. (**B**) DEGs downregulated under both osmotic and matrix stress. (**C**) DEGs uncorrelated across osmotic and matrix stress. For each pathway, the fraction of DEGs is noted (Rich Factor), the absolute number of genes upregulated in the pathway is the indicated by the symbol size (Count), and the adjusted P values are shaded (BH test; p-adjusted < 0.05, q < 0.2).

## DISCUSSION

Inoculation of plants with *Variovorax* relieves drought-induced stress (48, 54), increases biomass and yield (55, 56), and enhances nutrient uptake (57). The results described here show that distinct sets of DEGs arise from increasing water potential by changing osmolyte concentration and by changing matrix texture, which both occur as soils dry and *Variovorax* relieves plant stress. Across the two water potential parameters probed, ∼2.5-fold more DEGs were observed when changing the matrix compared with altering the osmolyte concentration. Although osmotic pressure changes resulted in a smaller number of DEGs compared with changes in matric potential, 76% of the DEGs observed when adding an osmolyte were also observed when changing the matrix. Furthermore, 67% of the shared DEGs presented the same direction of change. This observation identifies a set of DEGs that arise from increases in water potential, independent of the mechanism by which the water potential is changed. Prior transcriptomic studies have revealed shared effects of different osmolytes on transcription, include salts and organic solutes (23–28). Our study extends this shared response to gene expression changes arising from matrix stress, and it identifies the gene expression changes that are unique to water potential changes arising from growing cells in a textured material.

The increased complexity of the gene expression response across the matrix conditions could arise through multiple mechanisms. First, the matrices could affect the bioavailable concentration of chemicals in the mM9 growth medium. Prior studies have shown that soil matrices can differentially sorb apolar molecules, affecting the bioavailabile concentration of metabolites (58, 59), cell-cell signals (31), and macromolecules (60, 61). Second, ions in the growth medium could interact differently with the montmorillonite in M2 and the clay-sized quartz in Q2 because these materials differ in their cation exchange capacities (62). Third, each matrix may create distinct heterogeneous environments through the interaction with water, as hydration pockets in the different matrices may vary in their partitioning of cells and nutrients (63). In this way, matrices can have different effects on the extent to which microbes can access nutrients, whose diffusion is limited by the connectivity of hydration pockets (33). Finally, the matrices may not be perfectly matched in their pH, which can lead DEG pattern changes (64), even though the medium used for hydration of Q2 and M2 contains the same phosphate buffer. To understand how matrices and growth medium with different properties affect gene expression across a wider range of osmotic and matric potentials, future studies should examine DEGs across a wider range of artificial matrices that are hydrated using medium with defined compositions and osmotic potentials (31).

It is unclear how the trends observed might translate to other soil microbes within the context of their microbiomes, where cell-cell interactions also affect gene expression. A recent study that used a range of -omics tools to study how a reduced-complexity microbial consortium responds to changes in moisture and matrix by performing experiments in a glass bead porous medium amended with chitin (33). In this study, which used a model community containing *V. beijingensis* (33), the matrix influenced the microbial phenotypes observed, consistent with our findings with the Q2 and M2 soils having distinct textures. Another study investigating dynamic transcriptional changes, using a model community that includes *V. beijingensis*, observed a temporal shift in gene expression during chitin decomposition (65). Taken together, these studies show that to understand differences in osmotic and matrix stress more deeply, there will be a need to go beyond the static snapshot of gene expression in a single microbe and to examine how microbial communities respond dynamically to osmotic and matrix stress.

## ACKNOWLEDGEMENTS

This research was supported by a subcontract from the US Department of Energy (DOE), Office of Science, through the Genomic Science Program, Office of Biological and Environmental Research, under FWP 78814 at PNNL (to CAM and JJS). PNNL is a multi-program national laboratory operated by Battelle for the DOE under Contract DE-AC05-76RLO 1830. Support was also provided by the Office of Basic Energy Sciences of the U.S. Department of Energy grant DE-SC0014462 and by a National Science Foundation Research Traineeship Program 1828869 (to JJS).

## SUPPLEMENTAL MATERIAL

### Supplemental figures and tables

Figures S1 to S5

Tables S1 to S2

### Data set S1

Differential gene expression data

## DATA AVAILABILITY STATEMENT

All RNA-seq data has been submitted to the gene expression omnibus (GEO) under accession GSE299872. The code developed for this study can be accessed at https://github.com/SilbergLabRice/SeqSorcerer.

## REFERENCES

1. Milly PCD, Dunne KA, Vecchia AV. 2005. Global pattern of trends in streamflow and water availability in a changing climate. Nature 438:347–350.

2. Radolinski J, Vremec M, Wachter H, Birk S, Brüggemann N, Herndl M, Kahmen A, Nelson DB, Kübert A, Schaumberger A, Stumpp C, Tissink M, Werner C, Bahn M. 2025. Drought in a warmer, CO2 -rich climate restricts grassland water use and soil water mixing. Science 387:290–296.

3. Vereecken H, Amelung W, Bauke SL, Bogena H, Brüggemann N, Montzka C, Vanderborght J, Bechtold M, Blöschl G, Carminati A, Javaux M, Konings AG, Kusche J, Neuweiler I, Or D, Steele-Dunne S, Verhoef A, Young M, Zhang Y. 2022. Soil hydrology in the Earth system. Nat Rev Earth Environ 3:573–587.

4. Schimel JP. 2018. Life in Dry Soils: Effects of Drought on Soil Microbial Communities and Processes. Annu Rev Ecol Evol Syst 49:409–432.

5. Furtak K, Wolińska A. 2023. The impact of extreme weather events as a consequence of climate change on the soil moisture and on the quality of the soil environment and agriculture – A review. CATENA 231:107378.

6. Bardgett RD, Caruso T. 2020. Soil microbial community responses to climate extremes: resistance, resilience and transitions to alternative states. Phil Trans R Soc B 375:20190112.

7. Birch HF. 1958. The effect of soil drying on humus decomposition and nitrogen availability. Plant Soil 10:9–31.

8. Bottner P. 1985. Response of microbial biomass to alternate moist and dry conditions in a soil incubated with 14C- and 15N-labelled plant material. Soil Biology and Biochemistry 17:329–337.

9. Austin AT, Yahdjian L, Stark JM, Belnap J, Porporato A, Norton U, Ravetta DA, Schaeffer SM. 2004. Water pulses and biogeochemical cycles in arid and semiarid ecosystems. Oecologia 141:221–235.

10. Moyano FE, Manzoni S, Chenu C. 2013. Responses of soil heterotrophic respiration to moisture availability: An exploration of processes and models. Soil Biology and Biochemistry 59:72–85.

11. Manzoni S, Schimel JP, Porporato A. 2012. Responses of soil microbial communities to water stress: results from a meta-analysis. Ecology 93:930–938.

12. Van Gestel M, Merckx R, Vlassak K. 1993. Microbial biomass and activity in soils with fluctuating water contents. Geoderma 56:617–626.

13. Imminger S, Meier DV, Schintlmeister A, Legin A, Schnecker J, Richter A, Gillor O, Eichorst SA, Woebken D. 2024. Survival and rapid resuscitation permit limited productivity in desert microbial communities. Nat Commun 15:3056.

14. Singh S, Mayes MA, Kivlin SN, Jagadamma S. 2023. How the Birch effect differs in mechanisms and magnitudes due to soil texture. Soil Biology and Biochemistry 179:108973.

15. Warren CR. 2016. Do microbial osmolytes or extracellular depolymerisation products accumulate as soil dries? Soil Biology and Biochemistry 98:54–63.

16. Borken W, Matzner E. 2009. Reappraisal of drying and wetting effects on C and N mineralization and fluxes in soils. Global Change Biology 15:808–824.

17. Lado-Monserrat L, Lull C, Bautista I, Lidón A, Herrera R. 2014. Soil moisture increment as a controlling variable of the “Birch effect”. Interactions with the pre-wetting soil moisture and litter addition. Plant Soil 379:21–34.

18. Novick KA, Ficklin DL, Baldocchi D, Davis KJ, Ghezzehei TA, Konings AG, MacBean N, Raoult N, Scott RL, Shi Y, Sulman BN, Wood JD. 2022. Confronting the water potential information gap. Nat Geosci 15:158–164.

19. Luo S, Lu N, Zhang C, Likos W. 2022. Soil water potential: A historical perspective and recent breakthroughs. Vadose Zone Journal 21:e20203.

20. Harris RF. 2015. Effect of Water Potential on Microbial Growth and Activity, p. 23–95. In Parr, JF, Gardner, WR, Elliott, LF (eds.), SSSA Special Publications. Soil Science Society of America, Madison, WI, USA.

21. Naasko KI, Naylor D, Graham EB, Couvillion SP, Danczak R, Tolic N, Nicora C, Fransen S, Tao H, Hofmockel KS, Jansson JK. 2023. Influence of soil depth, irrigation, and plant genotype on the soil microbiome, metaphenome, and carbon chemistry. mBio 14:e01758–23.

22. Bremer E, Krämer R. 2019. Responses of Microorganisms to Osmotic Stress. Annu Rev Microbiol 73:313–334.

23. Yaakop AS, Chan K-G, Ee R, Lim YL, Lee S-K, Manan FA, Goh KM. 2016. Characterization of the mechanism of prolonged adaptation to osmotic stress of Jeotgalibacillus malaysiensis via genome and transcriptome sequencing analyses. Sci Rep 6:33660.

24. Wood JM, Bremer E, Csonka LN, Kraemer R, Poolman B, Van Der Heide T, Smith LT. 2001. Osmosensing and osmoregulatory compatible solute accumulation by bacteria. Comparative Biochemistry and Physiology Part A: Molecular & Integrative Physiology 130:437–460.

25. Imhoff JF, Rodriguez-Valera F. 1984. Betaine is the main compatible solute of halophilic eubacteria. J Bacteriol 160:478–479.

26. Kempf B, Bremer E. 1998. Uptake and synthesis of compatible solutes as microbial stress responses to high-osmolality environments. Archives of Microbiology 170:319–330.

27. Zhang Z-J, Chen S-H, Wang S-M, Luo H-Y. 2011. Characterization of extracellular polymeric substances from biofilm in the process of starting-up a partial nitrification process under salt stress. Appl Microbiol Biotechnol 89:1563–1571.

28. Upadhyay SK, Singh JS, Singh DP. 2011. Exopolysaccharide-Producing Plant Growth-Promoting Rhizobacteria Under Salinity Condition. Pedosphere 21:214– 222.

29. Fernandez-Illescas CP, Porporato A, Laio F, Rodriguez-Iturbe I. 2001. The ecohydrological role of soil texture in a water-limited ecosystem. Water Resources Research 37:2863–2872.

30. Rattray EAS, Prosser JI, Glover LA, Killham K. 1992. Matric potential in relation to survival and activity of a genetically modified microbial inoculum in soil. Soil Biology and Biochemistry 24:421–425.

31. Del Valle I, Gao X, Ghezzehei TA, Silberg JJ, Masiello CA. 2022. Artificial Soils Reveal Individual Factor Controls on Microbial Processes. mSystems 7:e00301–22.

32. Bello MO, Thion C, Gubry-Rangin C, Prosser JI. 2019. Differential sensitivity of ammonia oxidising archaea and bacteria to matric and osmotic potential. Soil Biology and Biochemistry 129:184–190.

33. Rodríguez-Ramos J, Sadler N, Zegeye EK, Farris Y, Purvine S, Couvillion S, Nelson WC, Hofmockel KS. 2025. Environmental matrix and moisture influence soil microbial phenotypes in a simplified porous media incubation. mSystems 10:e01616–24.

34. Kakumanu ML, Williams MA. 2014. Osmolyte dynamics and microbial communities vary in response to osmotic more than matric water deficit gradients in two soils. Soil Biology and Biochemistry 79:14–24.

35. Chowdhury N, Marschner P, Burns R. 2011. Response of microbial activity and community structure to decreasing soil osmotic and matric potential. Plant Soil 344:241–254.

36. Gao J, Sun Y, Xue J, Sun P, Yan H, Khan MS, Wang L, Zhang X, Sun J. 2020. Variovorax beijingensis sp. nov., a novel plant-associated bacterial species with plant growth-promoting potential isolated from different geographic regions of Beijing, China. Systematic and Applied Microbiology 43:126135.

37. McClure R, Farris Y, Danczak R, Nelson W, Song H-S, Kessell A, Lee J-Y, Couvillion S, Henry C, Jansson JK, Hofmockel KS. 2022. Interaction Networks Are Driven by Community-Responsive Phenotypes in a Chitin-Degrading Consortium of Soil Microbes. mSystems 7:e00372–22.

38. Wang Z, Gerstein M, Snyder M. 2009. RNA-Seq: a revolutionary tool for transcriptomics. Nat Rev Genet 10:57–63.

39. Marshall TJ, Holmes JW, Rose CW. 1996. Soil Physics, 3rd ed. Cambridge University Press. https://www.cambridge.org/core/product/identifier/9781139170673/type/book. Retrieved 19 June 2025.

40. Kim D, Paggi JM, Park C, Bennett C, Salzberg SL. 2019. Graph-based genome alignment and genotyping with HISAT2 and HISAT-genotype. Nat Biotechnol 37:907–915.

41. Li H, Handsaker B, Wysoker A, Fennell T, Ruan J, Homer N, Marth G, Abecasis G, Durbin R, 1000 Genome Project Data Processing Subgroup. 2009. The Sequence Alignment/Map format and SAMtools. Bioinformatics 25:2078–2079.

42. Liao Y, Smyth GK, Shi W. 2013. The Subread aligner: fast, accurate and scalable read mapping by seed-and-vote. Nucleic Acids Research 41:e108–e108.

43. Love MI, Huber W, Anders S. 2014. Moderated estimation of fold change and dispersion for RNA-seq data with DESeq2. Genome Biol 15:550.

44. Wickham H. 2016. ggplot2. Springer International Publishing, Cham. http://link.springer.com/10.1007/978-3-319-24277-4. Retrieved 12 June 2025.

45. Dixon P. 2003. VEGAN, a package of R functions for community ecology. J Vegetation Science 14:927–930.

46. Kanehisa M. 2000. KEGG: Kyoto Encyclopedia of Genes and Genomes. Nucleic Acids Research 28:27–30.

47. Wu T, Hu E, Xu S, Chen M, Guo P, Dai Z, Feng T, Zhou L, Tang W, Zhan L, Fu X, Liu S, Bo X, Yu G. 2021. clusterProfiler 4.0: A universal enrichment tool for interpreting omics data. The Innovation 2:100141.

48. Garcia Teijeiro R, Belimov AA, Dodd IC. 2020. Microbial inoculum development for ameliorating crop drought stress: A case study of Variovorax paradoxus 5C-2. New Biotechnology 56:103–113.

49. Fang J, Wei S, Gao Y, Zhang X, Cheng Y, Wang J, Ma J, Shi G, Bai L, Xie R, Zhao X, Ren Y, Lu Z. 2023. Character variation of root space microbial community composition in the response of drought-tolerant spring wheat to drought stress. Front Microbiol 14:1235708.

50. Trimble S. 2007. Encyclopedia of Water Science, Second Edition (Print Version). CRC Press. https://www.taylorfrancis.com/books/9781439846780. Retrieved 12 June 2025.

51. Cayley S, Lewis BA, Record MT. 1992. Origins of the osmoprotective properties of betaine and proline in Escherichia coli K-12. J Bacteriol 174:1586–1595.

52. Oren A. 1999. Bioenergetic Aspects of Halophilism. Microbiol Mol Biol Rev 63:334– 348.

53. Gupta A, Mishra R, Rai S, Bano A, Pathak N, Fujita M, Kumar M, Hasanuzzaman M. 2022. Mechanistic Insights of Plant Growth Promoting Bacteria Mediated Drought and Salt Stress Tolerance in Plants for Sustainable Agriculture. IJMS 23:3741.

54. Qi M, Berry JC, Veley KW, O’Connor L, Finkel OM, Salas-González I, Kuhs M, Jupe J, Holcomb E, Glavina Del Rio T, Creech C, Liu P, Tringe SG, Dangl JL, Schachtman DP, Bart RS. 2022. Identification of beneficial and detrimental bacteria impacting sorghum responses to drought using multi-scale and multi-system microbiome comparisons. The ISME Journal 16:1957–1969.

55. Natsagdorj O, Sakamoto H, Santiago DMO, Santiago CD, Orikasa Y, Okazaki K, Ikeda S, Ohwada T. 2019. Variovorax sp. Has an Optimum Cell Density to Fully Function as a Plant Growth Promoter. Microorganisms 7:82.

56. Belimov AA, Dodd IC, Hontzeas N, Theobald JC, Safronova VI, Davies WJ. 2009. Rhizosphere bacteria containing 1-aminocyclopropane-1-carboxylate deaminase increase yield of plants grown in drying soil via both local and systemic hormone signalling. New Phytologist 181:413–423.

57. Jiang F, Chen L, Belimov AA, Shaposhnikov AI, Gong F, Meng X, Hartung W, Jeschke DW, Davies WJ, Dodd IC. 2012. Multiple impacts of the plant growth-promoting rhizobacterium Variovorax paradoxus 5C-2 on nutrient and ABA relations of Pisum sativum. Journal of Experimental Botany 63:6421–6430.

58. Novak JM, Jayachandran K, Moorman TB, Weber JB. 2015. Sorption and Binding of Organic Compounds in Soils and Their Relation to Bioavailability, p. 13–31. In Skipper, HD, Turco, RF (eds.), SSSA Special Publications. Soil Science Society of America, American Society of Agronomy, and Crop Science Society of America, Madison, WI, USA.

59. Henriksen T, Svensmark B, Juhler RK. 2004. Degradation and Sorption of Metribuzin and Primary Metabolites in a Sandy Soil. J of Env Quality 33:619–627.

60. Demanèche S, Jocteur-Monrozier L, Quiquampoix H, Simonet P. 2001. Evaluation of Biological and Physical Protection against Nuclease Degradation of Clay-Bound Plasmid DNA. Appl Environ Microbiol 67:293–299.

61. G. P M. F, E. G, P. N. 2001. Effect of molecular characteristics of DNA on its adsorption and binding on homoionic montmorillonite and kaolinite. Biology and Fertility of Soils 33:402–409.

62. Dolcater DL, Lotse EG, Syers JK, Jackson ML. 1968. Cation Exchange Selectivity of Some Clay-Sized Minerals and Soil Materials. Soil Science Soc of Amer J 32:795–798.

63. Tecon R, Ebrahimi A, Kleyer H, Erev Levi S, Or D. 2018. Cell-to-cell bacterial interactions promoted by drier conditions on soil surfaces. Proc Natl Acad Sci USA 115:9791–9796.

64. Wilks JC, Kitko RD, Cleeton SH, Lee GE, Ugwu CS, Jones BD, BonDurant SS, Slonczewski JL. 2009. Acid and Base Stress and Transcriptomic Responses in *Bacillus subtilis*. Appl Environ Microbiol 75:981–990.

65. McClure R, Rivas-Ubach A, Hixson KK, Farris Y, Garcia M, Danczak R, Davison M, Paurus VL, Jansson JK. 2025. Multi-omics of a model bacterial consortium deciphers details of chitin decomposition in soil. mBio e00404–25.

