## Supporting Figures and Tables for "Effects of matric versus osmotic potential changes on *Variovorax beijingensis* transcription"

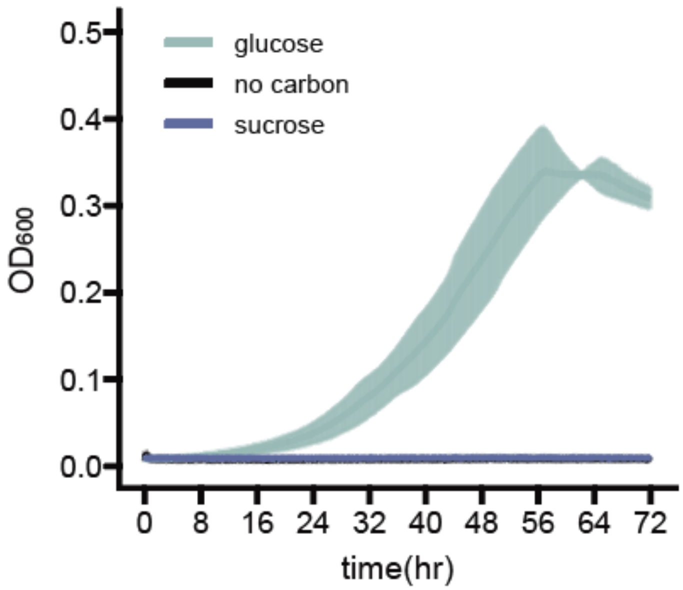


**Figure S1. *V. beijingensis* can use glucose but not sucrose as a carbon source.** Optical density ($OD_{600}$) of *V. beijingensis* grown in liquid M9 medium containing either glucose (*green*) and sucrose (*blue*). As a frame of reference cultures grown without a carbon source (*black*) are shown. Cultures were inoculated at a density of 0.05 and grown aerobically at 30°C while shaking at 108 rpm and 2.5 mm amplitude in a plate reader. For each experiment, 3 biological replicates are shown. The lines represent the average, while the shaded region represents ±1 standard deviations.

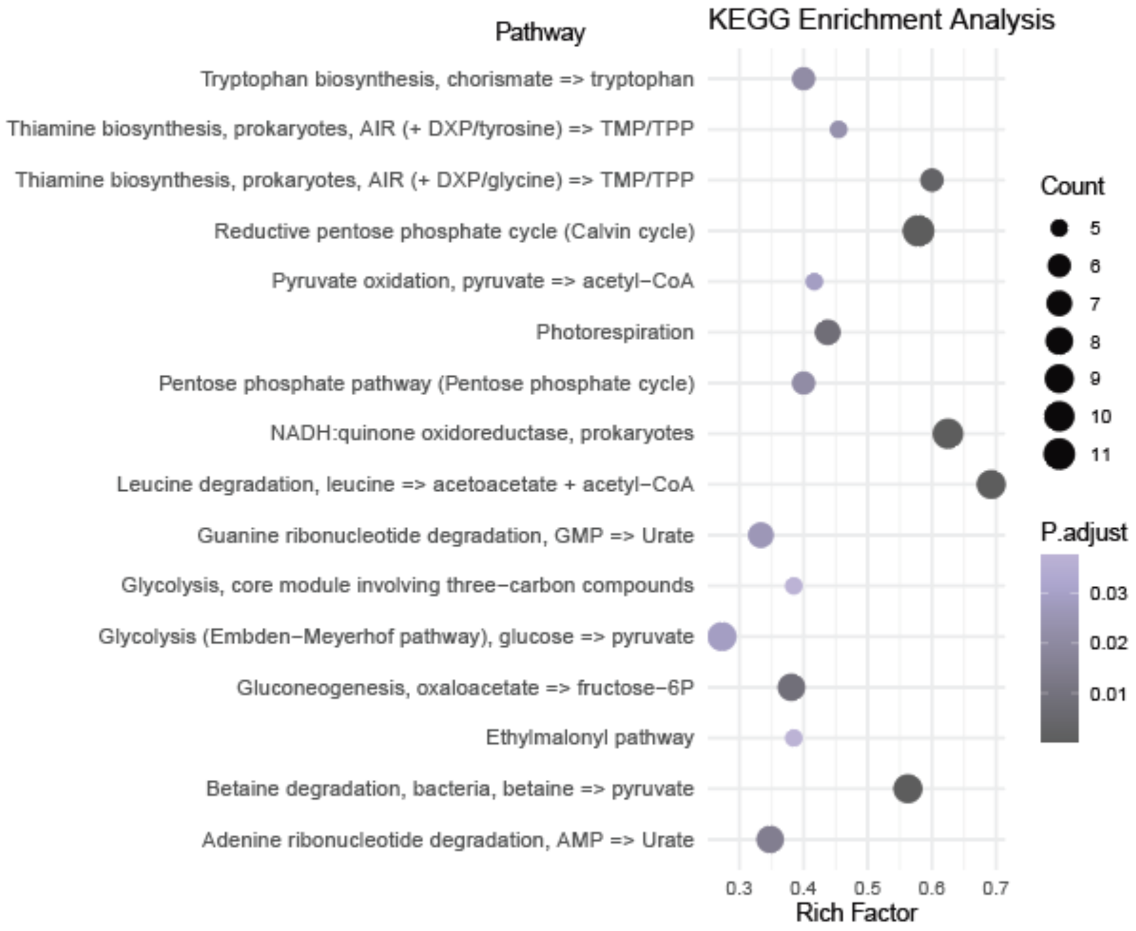


**Figure S2. KEGG enrichment analysis of genes upregulated in high osmotic potential compared to low potential.** For each pathway, the fraction of DEGs is noted (Rich Factor), the absolute number of genes upregulated in the pathway is the indicated by the symbol size (count), and the adjusted P values are shaded (BH test; p-adjusted < 0.05, q< 0.2).

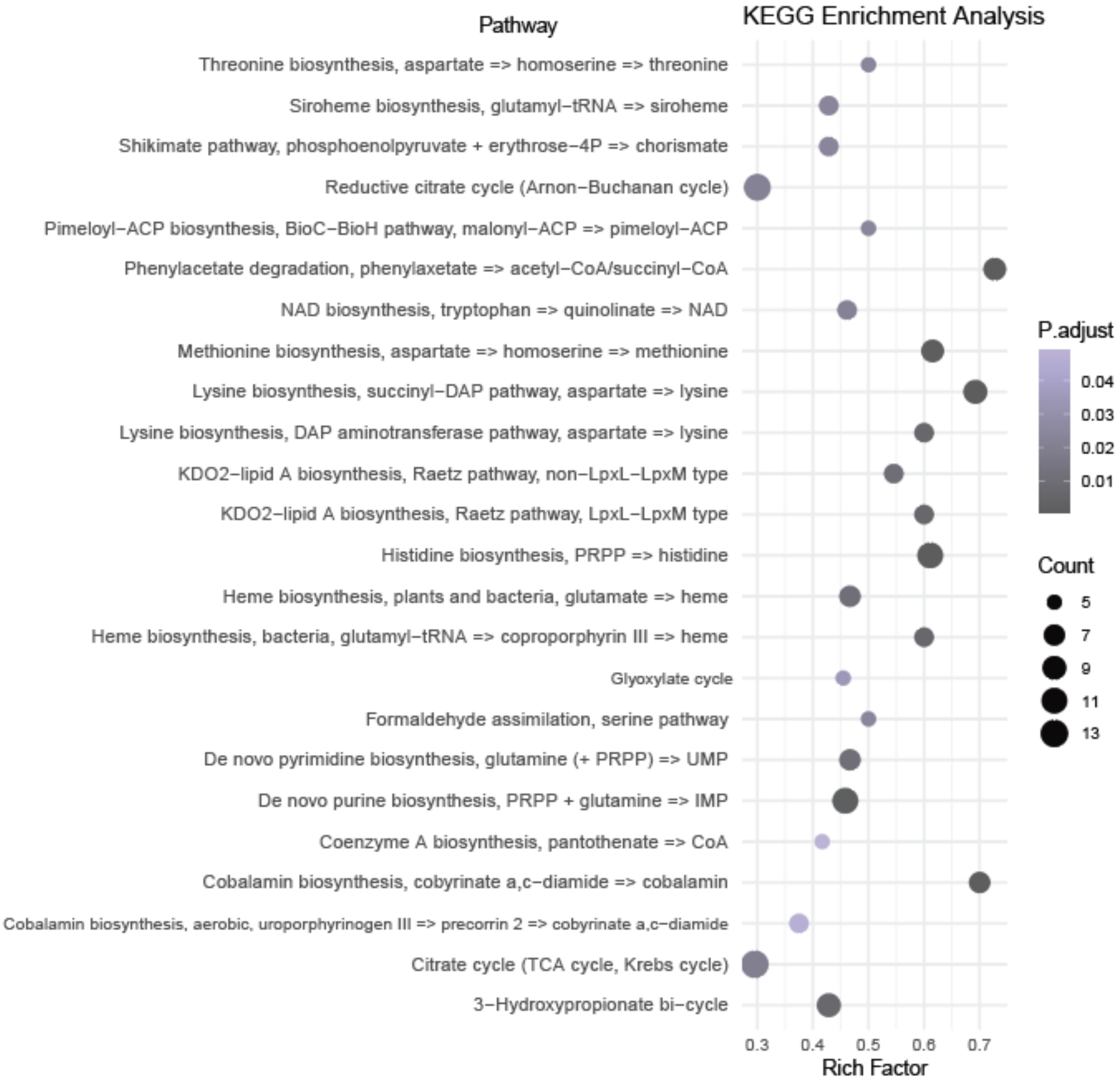


**Figure S3. KEGG enrichment analysis of down-regulated genes in high vs. low osmotic potential.** For each pathway, the fraction of DEGs is noted (Rich Factor), the absolute number of genes upregulated in the pathway is the indicated by the symbol size (count), and the adjusted P values are shaded (BH test; p-adjusted < 0.05, q< 0.2).

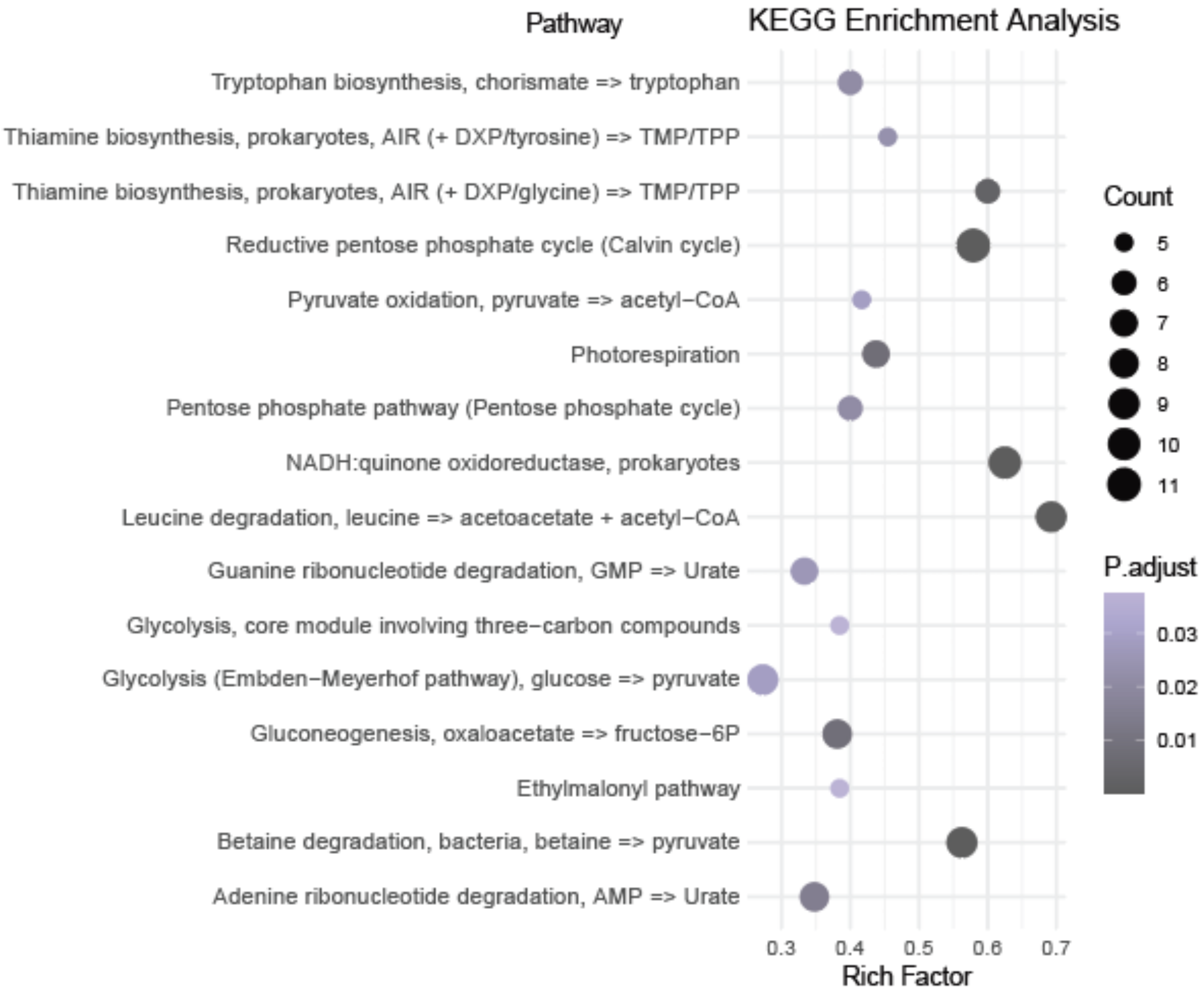


**Figure S4. KEGG enrichment analysis of upregulated genes in high vs. low matric potential.** For each pathway, the fraction of DEGs is noted (Rich Factor), the absolute number of genes upregulated in the pathway is the indicated by the symbol size (count), and the adjusted P values are shaded (BH test; p-adjusted < 0.05, q< 0.2).

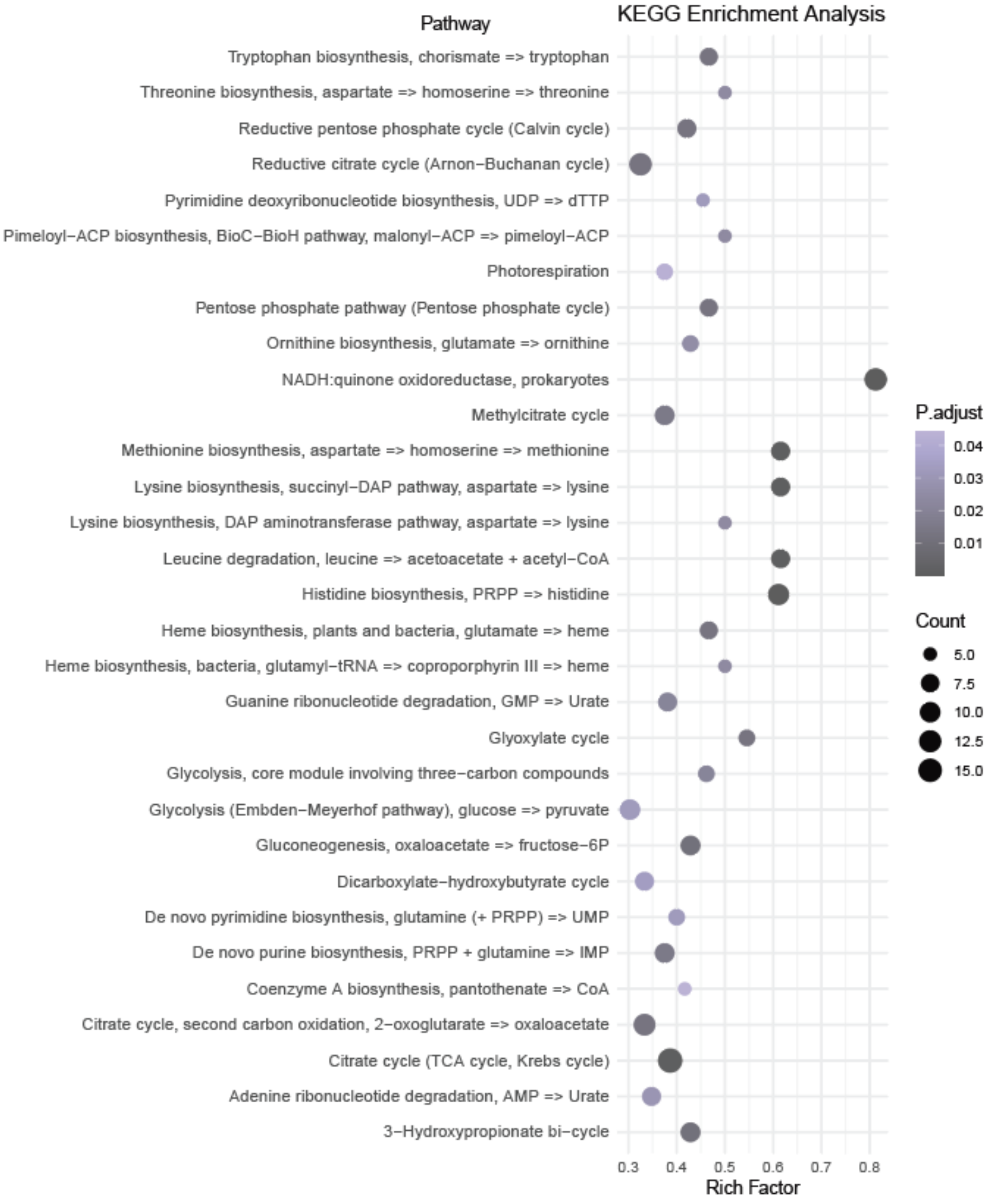


**Figure S5. KEGG enrichment analysis of down-regulated genes in high vs. low matric potential.** For each pathway, the fraction of DEGs is noted (Rich Factor), the absolute number of genes upregulated in the pathway is the indicated by the symbol size (count), and the adjusted P values are shaded (BH test; p-adjusted < 0.05, q< 0.2).

**Table S1. Pairwise Euclidean distance between conditions.** Based on the first two principal components calculated from the PCA, pairwise Euclidean distance was examined. The distance is a relative number, and bigger numbers represent greater variability between two groups. RNA sequencing data from measurements in Q2 and M2 are designated Low Ψm and High Ψm, respectively. Data from measurements in low and high osmolarity are Low Ψo and High Ψo, respectively.

| | **High Ψm** | **Low Ψm** | **High Ψo** | **Low Ψo** |
|---|---|---|---|---|
| **High Ψm** | 0 | 71.9462 | 71.9114 | 78.8553 |
| **Low Ψm** | 71.9462 | 0 | 52.94916 | 61.92812 |
| **High Ψo** | 71.9114 | 52.94916 | 0 | 9.587551 |
| **Low Ψo** | 78.8553 | 61.92812 | 9.587551 | 0 |

**Table S2. Permutational multivariate analysis of variance (PERMANOVA; 999 permutations) of RNA-seq data by condition (high osmotic-, low osmotic-, high matric-, and low matric-potential).** The first row “Model” shows statistical test performed by condition, and the second row “Residual” shows test performed by sample. Df represents degrees of freedom, SumSq represents sum of squares, $R^2$ shows % of total variation explained by the condition, F (F-statistics) is a ratio of inter-group to intra-group variation distance, and Pr(>F) represents *p*-value.

| | Df | SumSq | R2 | F | Pr(>F) |
|---|---|---|---|---|---|
| **Model** | 3 | 97123.49 | 0.822308 | 29.30894 | 0.001 |
| **Residual** | 19 | 20987.3 | 0.177692 | NA | NA |
| **Total** | 22 | 118110.8 | 1 | NA | NA |
